# *C. elegans* pharyngeal pumping provides a whole organism bio-assay to investigate anti-cholinesterase intoxication and antidotes

**DOI:** 10.1101/2020.06.03.131490

**Authors:** Patricia G. Izquierdo, Vincent O’Connor, Christopher Green, Lindy Holden-Dye, John Tattersall

**Author notes:** **Author for correspondence:** Patricia G. Izquierdo, School of Biological Sciences, Life Science Building 85, Room 3041, Highfield Campus, University of Southampton, Southampton United Kingdom, SO17 1BJ. **Research Highlights** - *C. elegans* pharyngeal pumping inhibition by organophosphates correlates with worm acetylcholinesterase inhibition by the anti-cholinesterases. - The recovery of the pharyngeal function in *C. elegans* in the presence of obidoxime is due to the recovery of the acetylcholinesterase function after anti-cholinesterase intoxication. - The pharyngeal neuromuscular function represents a quantitative bio-assay for investigation of anti-cholinesterase toxicity and recovery with excellent 3Rs potential.

## Abstract

Inhibition of acetylcholinesterase by either organophosphates or carbamates causes anti-cholinesterase poisoning. This arises through a wide range of neurotoxic effects triggered by the overstimulation of the cholinergic receptors at synapses and neuromuscular junctions. Without intervention, this poisoning can lead to profound toxic effects, including death, and the incomplete efficacy of the current treatments, particularly for oxime-insensitive agents, provokes the need to find better antidotes. Here we show how the non-parasitic nematode *Caenorhabditis elegans* offers an excellent tool for investigating the acetylcholinesterase intoxication. The *C. elegans* neuromuscular junctions show a high degree of molecular and functional conservation with the cholinergic transmission that operates in the autonomic, central and neuromuscular synapses in mammals. In fact, the anti-cholinesterase intoxication of the worm’s body wall neuromuscular junction has been unprecedented in understanding molecular determinants of cholinergic function in nematodes and other organisms. We extend the use of the model organism’s feeding behaviour as a tool to investigate carbamate and organophosphate mode of action. We show that inhibition of the cholinergic-dependent rhythmic pumping of the pharyngeal muscle correlates with the inhibition of the acetylcholinesterase activity caused by aldicarb, paraoxons and DFP exposure. Further, this bio-assay allows one to address oxime dependent reversal of cholinesterase inhibition in the context of whole organism recovery. Interestingly, the recovery of the pharyngeal function after such anti-cholinesterase poisoning represents a sensitive and easily quantifiable phenotype that is indicative of the spontaneous recovery or irreversible modification of the worm acetylcholinesterase after inhibition. These observations highlight the pharynx of *C. elegans* as a new tractable approach to explore anti-cholinesterase intoxication and recovery with the potential to resolve critical genetic determinants of these neurotoxins’ mode of action.

## 1. Introduction

Organophosphates and carbamates are potent acetylcholinesterase inhibitors (Colovic, Krstic et al. 2013, Tattersall 2018). This enzyme is key in terminating the cholinergic transmission that controls neuromuscular junction and important central synapse function (Koelle 1954, Massoulie, Pezzementi et al. 1993). This mode of action has led to the development of these compounds for widespread use as pesticides based on the central role of cholinergic transmission in the animal and plant parasitic life cycle (Takahashi and Hashizume 2014). This widespread use of anti-cholinesterases as pesticides has an associated human intoxication issue. At least two million cases of poisoning per year result in an estimated 200,000 deaths (Jeyaratnam 1990, Eddleston and Phillips 2004, Gunnell, Eddleston et al. 2007, Eddleston and Chowdhury 2016). Additionally acetylcholinesterase inhibitors with high human toxicity were developed as nerve agents for chemical warfare and terrorism (Colovic, Krstic et al. 2013, Worek, Wille et al. 2016).

The toxicological effect of organophosphates and carbamates is exerted through the covalent modification of acetylcholinesterase (AChE) (Colovic, Krstic et al. 2013, Tattersall 2018). The anti-cholinesterase drugs are orientated in the catalytic centre of the enzyme in a similar manner to acetylcholine (Dvir, Silman et al. 2010). When the molecule is positioned at the catalytic triad (Ser-Glu-His), the phosphorylation (OP) or carbamylation (carbamate) of the serine leads to inactivation of the AChE (Dvir, Silman et al. 2010). This inhibition results in the accumulation of the acetylcholine in the synaptic cleft causing the potential continued agonist activation of the two distinct classes of cholinergic receptors, muscarinic and nicotinic (Albuquerque, Deshpande et al. 1985). This overstimulation of the cholinergic target cells causes a wide range of neurotoxic effects. The first manifestations of the associated cholinergic syndrome cause autonomic disturbances including excessive sweating, lacrimation, salivation as well as cramps, bradycardia and miosis (Jokanovic and Kosanovic 2010, Tattersall 2018). Fatality occurs primarily due to disruption of the respiratory centres in the brain and/or transmission failure at the respiratory muscles (Jokanovic and Kosanovic 2010).

After enzyme inactivation, spontaneous reactivation occurs via hydrolysis of the bond created between the enzyme and the inhibitor molecule and enables the re-use of the AChE (Colovic, Krstic et al. 2013). This reversibility is important in managing recovery from intoxication. The rate at which it happens depends on the organophosphate or carbamate molecule and shows strong variation in the rate between distinct classes of anti-cholinesterase (Worek, Thiermann et al. 2004). However, the chemistry of the organophosphate attack is complicated by an ancillary reaction termed aging that leads to an irreversible inhibition in the OP-inhibited AChE (Wiener and Hoffman 2004, Colovic, Krstic et al. 2013). The dealkylation of any side chain of the conjugated OP creates a bond resistant to hydrolysis between the inhibitor and the catalytic serine (Li, Schopfer et al. 2007). It is a time-dependent reaction whose rate is extremely variable depending on the chemical structure of the intoxicating OP molecule (Worek, Thiermann et al. 2004).

Artificial ventilation is used to preserve breathing while the diaphragm neuromuscular junctions undergo a more long-term recovery involving hydrolysis and resynthesis of the otherwise dead molecule. This mitigation is supported by treatment with oximes, which are potent nucleophile molecules able to hydrolyse and reverse the AChE inhibition. In addition, the antidotes to intoxication are supported by low dose atropine and benzodiazepine treatment (Eddleston and Chowdhury 2016). Although this treatment has been used for the last 60 decades, it is deficient in several aspects (Buckley, Karalliedde et al. 2004). Firstly, atropine is a nonspecific competitive antagonist of muscarinic receptors, meaning that overstimulation of the target cell can still occur throughout the nicotinic receptors. Furthermore, there has not been a dose-response study to identify the optimal dose of atropine and the excessive administration might result in anti-muscarinic toxicity with fatal consequences (Eddleston, Buckley et al. 2004, Eddleston and Chowdhury 2016). Secondly, the success of reactivating AChE by oximes depends on which of the various OP molecules has produced the inhibition. For example, obidoxime seems to be more efficient for reactivating AChE after the inhibition of OP pesticides but not nerve agents. The efficiency of 2-pralidoxime is demonstrated after the inhibition of AChE with sarin or VX but not by soman or tabun. Lastly, there is not any reactivator able to recover the AChE activity after the aging reaction (Worek, Thiermann et al. 2004).

The limitations of the current treatment, poor health condition of the surviving victims and the fatalities reported have become a major public health concern (Jeyaratnam 1990, Konradsen 2007). In this scenario, identifying model organisms, which replicate the biological manifestation of OP toxicity, is key to develop alternative strategies. Mammal animal models have been used to address this situation, with species ranging from small rodents to large mammals, including non-human primates (Pereira, Aracava et al. 2014). The signs and LD50 values of anti-cholinesterase poisoning in these models are well correlated to the IC50 of AChE inhibition in both brain and blood samples (Sivam, Hoskins et al. 1984, Fawcett, Aracava et al. 2009). However, since the development of the current treatment, between 1950s and 1960s, it has not been significantly improved. Taking into consideration this fact as well as the 3Rs principles for animal research (Prescott and Lidster 2017, Balls and Combes 2019), the genetically tractable model organism *C. elegans* is proposed in this study. It has been widely used in neurotoxicological studies including organophosphates (Cole, Anderson et al. 2004, Melstrom and Williams 2007, Rajini, Melstrom et al. 2008, Jadhav and Rajini 2009, Lewis, Szilagyi et al. 2009, Vinuela, Snoek et al. 2010, McVey, Mink et al. 2012, Leelaja and Rajini 2013, Lewis, Gehman et al. 2013). This is advantaged by highly conserved molecular pathways between the nematode and humans. There is a rich cholinergic signalling network in which acetylcholine controls the worm’s nervous system and is essential for neuromuscular transmission (Rand 2007, Pereira, Kratsios et al. 2015). The cholinergic neuromuscular transmission, which excites distinct muscles, underpins biologically critical functions such as locomotion, egg-laying and the feeding behaviour (Rand 2007, McVey, Mink et al. 2012). As in mammals, acetylcholinesterase is key in terminating the cholinergic signal to prevent hyperstimulation. The three *C. elegans* acetylcholinesterases are orthologous to the three human acetylcholinesterase isoforms (Arpagaus, Combes et al. 1998, Combes, Fedon et al. 2000, Selkirk, Lazari et al. 2005). In particular, the catalytic centre of the nematode enzyme is highly conserved to mammals and harbours the key amino acids involved in the inhibition and aging reactions (Combes, Fedon et al. 2000).

*C. elegans* exposed to anti-cholinesterases exhibits easily scored defects in behaviours (McVey, Mink et al. 2012). These include hypercontraction of body wall muscles that results in the paralysis of the worms (Cole, Anderson et al. 2004, Rajini, Melstrom et al. 2008, McVey, Mink et al. 2012). In fact, toxicity ranks for investigated anti-cholinesterases are comparable in their inhibition to relative potencies identified in mammalian models, consistent with the highly conserved catalytic site (Cole, Anderson et al. 2004, Rajini, Melstrom et al. 2008).

We have investigated how anti-cholinesterases act on the high rate of pharyngeal pumping that worms use to feed on bacteria (Avery 1993, Avery and Shtonda 2003, Niacaris and Avery 2003). Here, we show that whole organism measurement of pharyngeal movements represents a sensitive phenotype that allows us to use it as a bio-assay for the whole organism effects of OP intoxication. Furthermore, the inhibition of nematode acetylcholinesterases was better correlated to the inhibition of the pharyngeal pumping than to the paralysis of the body wall muscles. We validated the pharyngeal pumping as a tool to probe spontaneous recovery as well as the reversible and irreversible inhibition associated with aging. This was confirmed by biochemical analysis of the nematode acetylcholinesterase activity. Thus, the pharynx offers a powerful bio-assay to investigate mode of action and approaches by which chemical mitigation can be used to treat intoxication. The possibility to resolve genetic determinants that might act beyond the primary mode of action of the organophosphate in *C. elegans* suggests the organization of pharyngeal pumping might provide a route to allow novel understanding of these important neurotoxins.

## 2. Materials and Methods

### 2.1. C. elegans maintenance

All the experiments were performed using N2 Wild-type *C. elegans* strain obtained from Caenorhabditis Genetics Center (https://cgc.umn.edu/) and maintained under standard conditions (Brenner 1974). Briefly, nematodes were growth at 20°C on Nematode Growth Medium (NGM) agar plates seeded with *E. coli* OP50 as source of food.

### 2.2. Drug stocks

Carbamate (aldicarb) and organophosphates (paraoxon-ethyl, paraoxon-methyl and DFP) were acquired from Merck and dissolved in 70% ethanol and 100% DMSO, respectively. The oximes, obidoxime and 2-pralidoxime, were provided by DSTL Porton Down (UK) and dissolved in distilled autoclaved water. The drug stocks were kept at 4°C, as manufacturer recommended temperature, in a locked cabinet according with standard security protocols. Dissolved compounds were used within one month or discarded.

Acetylthiocholine iodide (ATCh) and 5,5’-dithio-bis-2-nitrobenzoic acid (DTNB) were obtained from Merck (https://www.sigmaaldrich.com/united-kingdom.html) and dissolved in phosphate buffer 0.1 M pH7.4 directly before use.

### 2.3. Behavioural assays

All behavioural experiments were performed on a standard developmental stage: young hermaphrodite adults (L4 + 1 day) at room temperature (20°C). Worms were allowed to develop from eggs at 20°C through the larval stages L1, L2, L3. Worms were viewed under a Nikon SMZ800 binocular zoom microscope and were recognized as L4 by the temporary appearance (8 hours window) of the vulva saddle. These worms were selected the day before the experiment and placed on fresh OP50 seeded plates. They were used 16-24 hours after as L4 +1.

Anti-cholinesterase containing plates were prepared the day before of each experiment by adding an aliquot of a concentrated stock in the melted NGM, tempered after heating to approximately 60°C. 50 μl of OP50 *E. coli* bacteria one OD_600 nm_ was dropped on the plate when the media was solidified. After 1 hour in the fume hood, the dried bacterial plates were sealed and kept in dark at 4°C until next day. Plates were left at room temperature for at least 30 min before starting the experiment. There was no observable change in the bacterial lawn of anti-cholinesterase-containing and control plates, therefore no effect of the anti-cholinesterase on the *E. coli* growth was discernible (Kudelska, Lewis et al. 2018). The final concentration of vehicle in the behavioural assay was 0.07% ethanol for aldicarb-containing plates and 0.1% of DMSO for organophosphate-containing plates. Control plates contained the same concentration of vehicle than assay plates. Neither vehicle concentrations alone had any effect in the phenotypes tested.

Nematodes were picked onto the bacterial lawn 10 minutes before starting observations. Worms that left the patch of food during the experiment were picked back to the bacterial lawn. They had to be on the lawn for at least 10 minutes before the start of observations to be counted (Dalliere, Bhatla et al. 2016).

### 2.4. Intoxication with aldicarb

Experiments were performed in 6-well plates containing a final NGM volume of 3 ml. Aldicarb was added to the assay at final concentration between 2 μM and 500 μM. Vehicle control plates were used as control. Young adult worms were placed on aldicarb and non-aldicarb containing plates where the pharyngeal pumping, percentage of paralysed worms and body length was scored at indicated times.

Paralysis and body length were scored as previously described (Mahoney, Luo et al. 2006, Mulcahy, Holden-Dye et al. 2013). Briefly, nematodes were picked onto the bacterial lawn containing either aldicarb or vehicle control. Paralysis was scored by quantifying the number of animals not moving out the total of worms on the plate at indicated times. These snapshots involved scoring for 30 secs. Nematodes were considered paralysed when no movement was detected after prodding three times with a platinum wire (Mahoney, Luo et al. 2006). To measure body length, images of the worms were taken at the specified times. These images of the nematodes were binarized and skeletonized using ImageJ software. The length of the skeleton was used to determine the body length of the nematodes (Mulcahy, Holden-Dye et al. 2013).

Pharyngeal pumping on food in the presence of aldicarb was scored at indicated times after transferring worms to aldicarb or vehicle control plates (10 min, 1 h, 2 h, 6 h, and 24 h after picking onto assay plates). Pumping was quantified by counting the number of grinder movements observed under binocular microscope. The pump rate was quantified for a minimum of 3 minutes per worm at each time point and the mean was used as pumps per minute.

The estimates of IC50 were made by measuring the body wall and pharyngeal function at varying drug concentrations after 24 hours incubation relative to worms placed on drug free vehicle control plates. The percentages of inhibition relative to these controls were used to estimate IC50.

### 2.5. Intoxication with the organophosphates paraoxon-ethyl, paraoxon-methyl and DFP

For pharyngeal intoxication assays with organophosphates, the same procedures were utilized as indicated in section 2.4., with the exception of DFP. This organophosphate equilibrates across the individual wells of the 6-well culture plates. This was evidenced by the inhibition of pumping and paralysis of worms placed on non-DFP wells adjacent to DFP laced agar (data not shown). This potent cross-contamination by DFP concentrations precluded the use of 6-well plates. Therefore, 9 cm Petri dishes were used containing a final volume of 20 ml NGM. DFP was added to the melted NGM as mentioned above to obtain the indicated final concentrations between 2 μM and 500 μM. Non-DFP containing plates were used as control. After solidification, 200 μl of *E. coli* OP50 OD_600 nm_ = 1 was spread evenly over the complete surface of the NGM. This full food coverage was needed to mitigate the potent drive for worms to leave food that was particularly strong in the case of the DFP treatments (data not shown). Seeded plates were incubated for 1 h and then they were kept until the next day as mentioned in section 2.3.

### 2.6. Recovery from organophosphate intoxication

To study the recovery of pharyngeal pumping after organophosphate intoxication, L4 + 1 worms were intoxicated on organophosphate-containing plates for 24 hours. The intoxicating concentration was calculated based on estimating the lowest concentration that gave the maximal inhibition of pumping after 24 hours of exposure. After incubation on organophosphate laced plates for 24 h, the nematodes were transferred onto either non-drug containing plates or oxime-containing plates. From here, the recovery from full inhibition was measured by recording the pump rate at indicated times after being placed on no-drug or oxime plates. Oxime plates were poured, seeded with OP50 and stored using the protocol mentioned in section 2.3. Neither obidoxime nor pralidoxime alone had an effect on the pharyngeal pumping rate at concentrations between 0.5 mM and 2 mM (Supplementary figure 1).

### 2.7. Biochemical assays

Total worm homogenates were generated from synchronized L4/adult worms. For this, 12 gravid worms were maintained for 4 h on OP50 seeded 5.5 cm plates, in which time they accumulated freshly laid eggs. The adult worms were removed and plates were incubated 3 days at room temperature. This generated approximately 250 age-synchronized L4/adults on bacteria depleted plates. Worms from a minimum of 40 plates (approx. 10,000 worms) were harvested and washed three times with 0.1 M phosphate buffer pH7.4 in order to remove all the remaining bacteria. Nematodes were transferred to a glass homogenizer and incubated for 30 min on ice with a final concentration of 0.15% of n-octyl-glucoside as detergent to permeabilize the cuticle and release cellular content (Blaxter 1993). The n-octyl-glucoside did not alter the acetylcholinesterase activity (data not shown). Mouse brain homogenate was used in parallel to validate the acetylcholinesterase activity quantification protocols in *C. elegans* and compare with previously published data. To generate mouse forebrain homogenate, freshly dissected tissue was homogenized in 10 volumes of phosphate buffer (w/v). This was kindly provided by Aleksandra Pitera (Southampton University, UK). Worm/mouse protein homogenate was stored at −80°C until use when they were defrosted on ice.

Acetylcholinesterase activity was measured using a modified colorimetric Ellman’s assay (Ellman, Courtney et al. 1961). The assay mixture contained 0.2 mg/ml of worm/mouse homogenate comprising the AChE enzyme, 0.48 mM acetylthiocholine (ATCh) as substrate and 0.32 mM 5,5-dithio-bis-(2-nitrobenzoic acid) (DTNB) as chromophore in a final volume of 200 μl of 0.1 M phosphate buffer pH7.4. The increase in absorbance at 410 nm was measured at 1 min intervals for 15 min at room temperature using a FLUOstar Optima microplate reader (BMG Labtech). The change in absorbance against time due to the production of 5-thio-2-nitro-benzoic acid and its extinction coefficient was utilized to calculate the acetylcholinesterase activity (μmoles/min). This was normalized to the protein content of worm/mouse homogenate determined by standard Bradford protocol (Bradford 1976). The enzyme activity in the homogenate was expressed as μmoles/min/mg protein.

### 2.8. Acetylcholinesterase activity after whole worm aldicarb intoxication

To estimate the acetylcholinesterase activity after aldicarb intoxication, nematodes were synchronized as in section 2.7. When worms reached the L4/adult stage, an aliquot of 12 μl aldicarb stock was added to the worm-containing plates (12 ml) to generate the indicated final aldicarb concentration of 50 μM or 500 μM. Control plates were made by adding 12 μl of 70% ethanol. A minimum of 40 plates were used per condition. After 24 hours of incubation at room temperature the control and aldicarb intoxicated worms were harvested, washed and treated to generate the worm homogenate. Nematodes were kept on ice during the whole process to prevent recovery of the acetylcholinesterase activity through reversibility of the reaction. Acetylcholinesterase activity assays were carried out directly after the worm protein extraction.

### 2.9. Acetylcholinesterase activity of worm/mouse brain homogenate after inhibition by organophosphates

A stock solution of organophosphate was appropriately diluted in 0.1 M phosphate buffer pH7.4 just before the experiment to ensure a low concentration of DMSO in the final dilution for acetylcholinesterase activity quantification. The final concentration of vehicle in the biochemical assay was 0.000025%.

Worm/mouse brain homogenate in phosphate buffer (0.1 M pH7.4) as described above was placed in a 96-well plate. At 0 min, 15 min, 30 min, 35 min, 40 min, 43 min, 44 min, 44.3 min, 44.6 min and 45 min, these volumes were supplemented with organophosphate to the indicated final concentration in 50 μl final volume. This incubation contained 120 μg of worm/mouse protein and 1 μM organophosphate. After 45 min, acetylcholinesterase activity assay was scored for all the samples as described above (section 2.7) by addition of DNTB and acetylthiocholine in a final volume of 200 μl with 0.1 M phosphate buffer pH7.4. This gave a time series in which the time of incubation with the organophosphate was 20 s, 40 s, 1 min, 2 min, 5 min, 10 min, 15 min, 30 min and 45 min. An acetylcholinesterase assay without the presence of the OP was used as control.

Either single or double decay exponential curve was fitted to these time courses to determine the best-fit mode of inhibition of acetylcholinesterase by each organophosphate.

### 2.10. Acetylcholinesterase reactivation after inhibition with organophosphate drugs

Organophosphate-inhibited acetylcholinesterase was prepared by incubating 2.4 mg/ml of worm/mouse homogenate with 2 μM of any organophosphate in a final volume of 320 μl for 1 hour at room temperature. The mixture was centrifuged at 14000 rpm for 30 min and the supernatant was discarded to remove the excess of organophosphate. The pellet containing the inhibited acetylcholinesterase was resuspended with 320 μl of phosphate buffer (0.1 M pH7.4) and maintained at room temperature. For control, non-exposed 2.4 mg/ml worm/mouse protein was used in parallel. Following organophosphate removal, aliquots were taken at subsequent time intervals to determine acetylcholinesterase activity after incubating the mixture 1 min in the presence or absence of 100 μM obidoxime (Worek, Thiermann et al. 2004). The concentration of obidoxime used was previously described to research spontaneous recovery and aging reaction in human acetylcholinesterase inhibited by paraoxon-ethyl, paraoxon-methyl or DFP (Worek, Thiermann et al. 2004).

Acetylcholinesterase activity that does not recover upon incubation with obidoxime is considered to have arisen from a dealkylation reaction that ages the modified enzyme (Worek, Thiermann et al. 2004). Mouse brain acetylcholinesterase was used in order to validate the protocol and compare the nature of organophosphate mode of action in the two organisms (Kardos and Sultatos 2000).

### 2.11. Statistical analysis

Data were analysed using GraphPad Prism 7 and are given as mean ± SEM. Statistical significance was assessed using two-way ANOVA followed by post hoc analysis with Bonferroni corrections when applicable. Bonferroni corrections were selected to avoid false positives. The sample size N of each experiment is specified in the figure.

## 3. Results

### 3.1. Quantifying anti-cholinesterase induced changes in cholinergic neuromuscular function with whole organism behaviour

We first investigated distinct behaviours that are underpinned by cholinergic neuromuscular junction function in *C. elegans*. This identified that locomotion/paralysis, contraction mediated shrinkage of body length and the rate of pharyngeal pumping showed a clear concentration-time dependent inhibition with respect to this class of anti-cholinesterase. The carbamate aldicarb was used as representative of the acetylcholinesterase inhibitors. Similar to organophosphates, aldicarb binds and inhibits acetylcholinesterase, resulting in the increase of acetylcholine concentration at the neuromuscular junctions (Colovic, Krstic et al. 2013). This produces an overstimulation of the cholinergic receptors that leads to the hypercontraction of the muscle cells at different neuromuscular junctions in *C. elegans* (McVey, Mink et al. 2012). The aldicarb-induced hypercontraction of the body wall muscles elicited both paralysis and decrease of the body length of the nematodes (Fig. 1). However, the lowest concentrations of aldicarb tested (2 μM, 10 μM and 50 μM) failed to paralyse the worms, even though the nematodes incubated on 50 μM aldicarb plates for 24 hours looked uncoordinated and were significantly shorter than control nematodes (Fig. 1).

**Figure 1.**
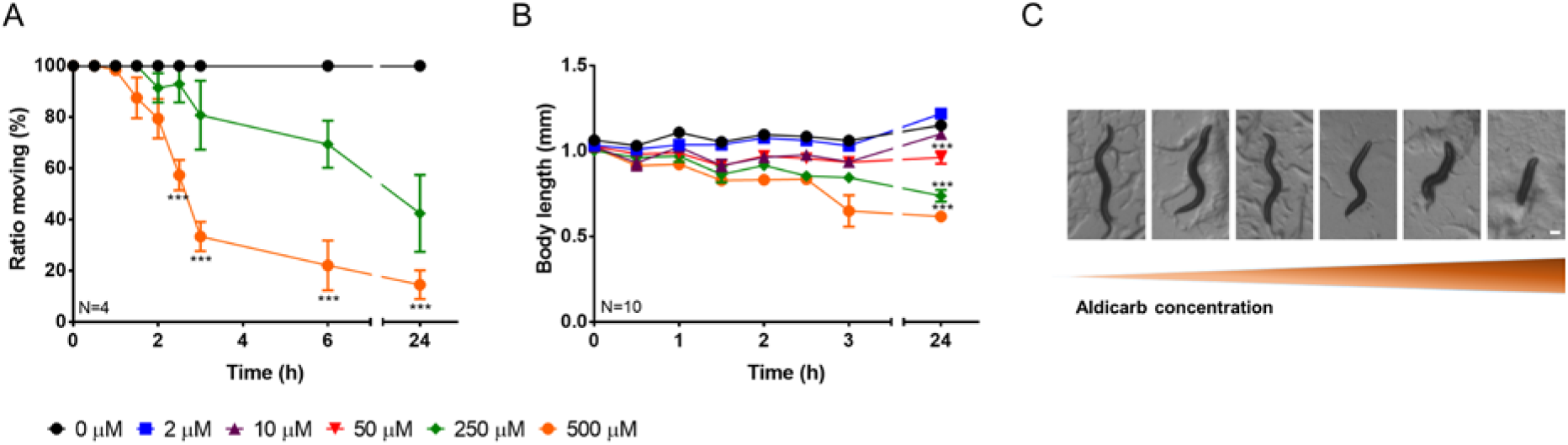
Nematodes exposed to aldicarb exhibited paralysis and hypercontraction of body wall muscles. A) The number of synchronized L4 + 1 nematodes moving as percentage the total worms on the plate was scored at different times in the face of a range of concentrations of aldicarb. The ratio moving of nematodes exposed to 2 μM, 10 μM and 50 μM was identical to the control. Data are shown as mean ± SEM of four different experiments. B) The body length of synchronized L4 + 1 nematodes exposed to different concentrations of aldicarb plates was scored by taking micrographs at different times of incubation and the length quantified. Data are shown as mean ± SEM of the length of ten worms in five different experiments. C) Body length of nematodes incubated on different concentrations of aldicarb plates for 24 hours. From the left to the right the concentration of aldicarb was: 0 μM (control), 2 μM, 10 μM, 50 μM, 250 μM and 500 μM. Scale bar represents 100 μm. *p<0.05; **p<0.01; ***p<0.001 by two-way ANOVA test.

In a similar way, the aldicarb treatment caused a dose-dependent inhibition of pharyngeal pumping. (Fig. 2). This paralysis is likely mediated by the elevated acetylcholine associated with cholinesterase inhibition. The consequence is the hyperstimulation of the radial muscle contractions that underpin rhythmic pumping mediated by a pacemaker cholinergic transmission (Avery 1993, Avery and Shtonda 2003, Niacaris and Avery 2003). This notion is supported by experiments in which the isolated pharynx is exposed to physiological Dent’s solution with or without 5 μM aldicarb for 5 min (Fig. 2B). In the carbamate-treated isolated pharynx, we observed the predicted hypercontraction of the pharyngeal radial muscle evidence by the sustained opening of the pharyngeal lumen (Fig. 2B). In contrast to body wall muscles, the whole organism paralysis of the pharyngeal muscle was observed with the lowest concentrations tested within 6 hours of incubation on the aldicarb-containing plate and from the first hour of intoxication onto 50 μM plates.

**Figure 2.**
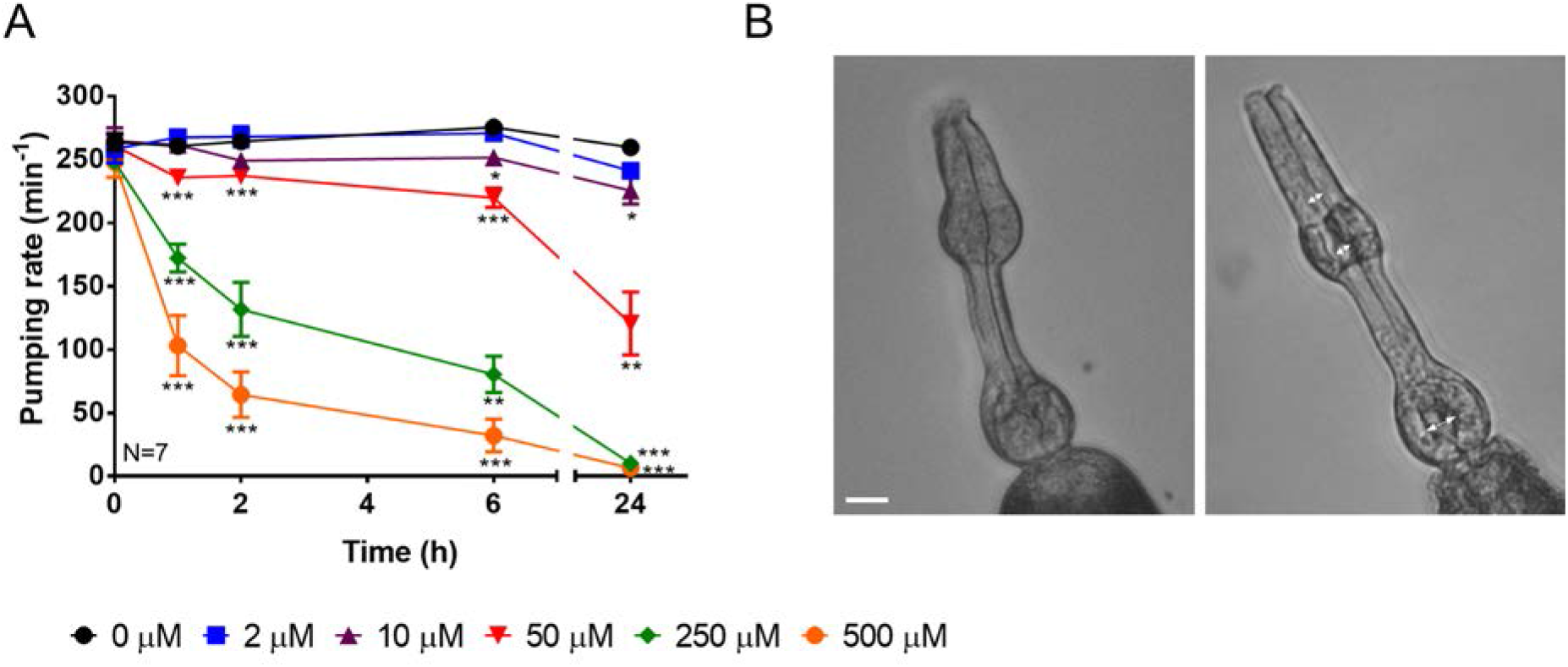
Pharyngeal pumping of *C. elegans* exposed to aldicarb exhibited a gradual concentration-time dependent paralysis due to the hypercontraction of the radial muscles in the pharynx. A) Pharyngeal pumping rate per minute was quantified at different end-point times for synchronized L4 + 1 nematodes exposed to a range of concentration of aldicarb plates. An increased concentration-dependent response over the time is observed. Data are shown as mean ± SEM of the pumping rate of 7 worms in four different experiments. B) Isolated pharynx of *C. elegans* were exposed to Dent’s solution as control (left panel) or 5 μM of aldicarb (right panel). The hypercontraction of the radial muscles caused the opening of the pharyngeal lumen (indicated by the white arrows). Scale bar represents 1 μm. *p<0.05; **p<0.01; ***p<0.001 by two-way ANOVA test.

The IC50 values calculated after 24 hours of intoxication indicated that pharyngeal pumping rate and body length were the most sensitive behaviours to anti-cholinesterase intoxication, being 5 fold lower than paralysis IC50 value (Fig. 3). Moreover, the pharyngeal pumping rate discriminated the incremental effect of increasing concentrations of cholinesterase inhibition. Furthermore, this whole organism bio-assay of cholinergic neuromuscular junction was more sensitive than body length in resolving the low concentration as well as discerning anti-cholinergic effects at shorter incubation times.

**Figure 3.**
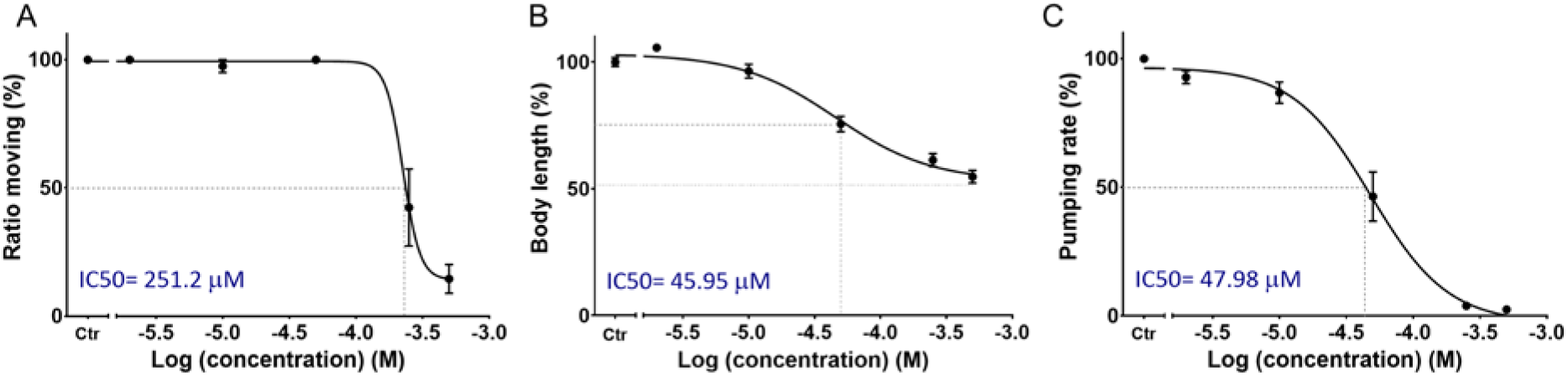
Aldicarb concentration-dependent sensitivity in cholinergic neuromuscular junction dependent behaviours. A) The percentage of ratio moving corresponds to the number of worms moving out of the total number of worms on the plates after 24 hours of intoxication. B) Body length was expressed as percentage of the unexposed body length after 24 hours of incubation. C) Pharyngeal pump rate was expressed as percentage of the pharyngeal pumping of unexposed nematodes after 24 hours of incubation. Data are shown as mean ± SEM. *p<0.05; **p<0.01; ***p<0.001 by two-way ANOVA test.

### 3.2. *C. elegans* acetylcholinesterase activity is reduced by the presence of aldicarb

In order to demonstrate that aldicarb inhibits acetylcholinesterase activity in the treated *C. elegans*, intoxicated worms on 50 μM and 500 μM aldicarb plates were harvested and homogenized. The acetylcholinesterase activity in the homogenates was measured and compared with non-treated animals (Fig. 4A). Nematodes intoxicated onto aldicarb plates for 24 hours exhibited a reduction in the acetylcholinesterase activity that was dependent on the inhibitor concentration (Fig. 4A). Interestingly, the inhibition of enzyme activity was greater than the reduction in any of the neuromuscular junction dependent phenotypes investigated. The nematodes incubated on 50 μM aldicarb displayed 15% of acetylcholinesterase activity found in control homogenates (Fig. 4A). In contrast, locomotion, body length and pharyngeal pumping were 100%, 83% and 53%, respectively, compared to the corresponding control (Fig. 4B, 4C, 4D). Similarly, the acetylcholinesterase activity of nematodes after 24 hours of incubation on 500 μM aldicarb plates was 3.4% while the ratio of worms moving, body length and pumping rate was 14.6%, 53.9% and 2.5%, respectively, compared to the corresponding control of non-treated worms (Fig. 4). These data indicate the high safety factor associated with cholinesterase function in which low levels of acetylcholinesterase activity can maintain the behavioural function in addition with a large reserve of acetylcholinesterase available to replace the organophosphate inhibited. This is consistent with previous data from mammal models in which the function was maintained despite profound enzyme inhibition (Wolthuis, Groen et al. 1995).

**Figure 4.**
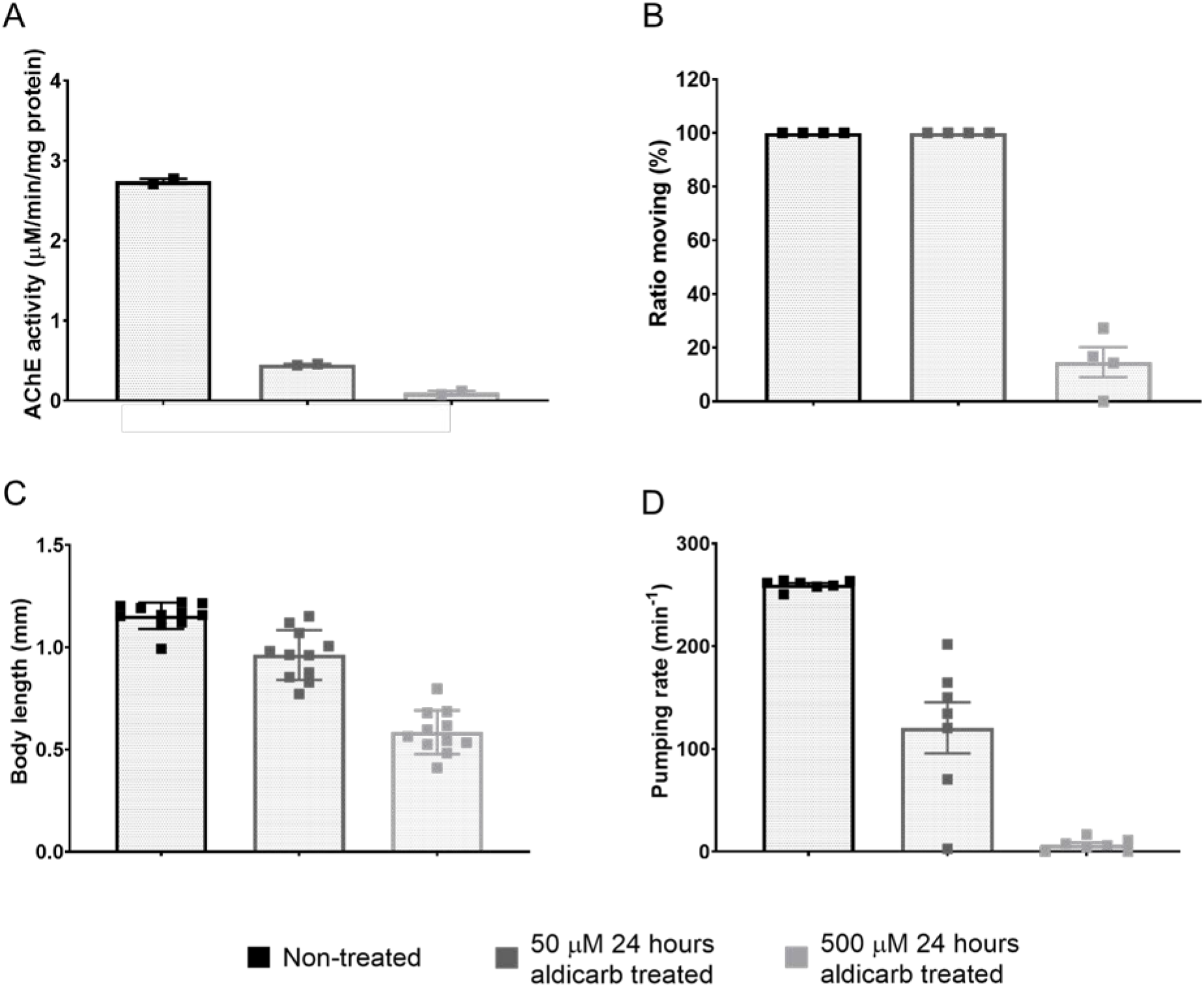
*C. elegans* acetylcholinesterase activity associated with reduced pharyngeal pumping rate and motility behaviours after 24 hours of intoxication. A) *C. elegans* acetylcholinesterase activity associated with homogenates from synchronized L4/adult worms isolated after 24 hours of incubation onto empty, 50 μM and 500 μM aldicarb plates. Treated worms were homogenized and enzyme activity measured using a modified Ellman’s assay. Data are shown as mean ± SEM of two independent experiments. B) The ratio moving was scored as the animals moving after 24 hours of exposure onto empty, 50 μM and 500 μM aldicarb plates as percentage of the total worms on the corresponding plate. Data are shown as mean ± SEM of four independent experiments. C) Body length of L4 + 1 nematodes was scored after 24 hours exposed to 50 μM, 500 μM aldicarb or unexposed. Data are shown as mean ± SEM of the length of 10 worms in five independent experiments. D) Pharyngeal pumping rate of unexposed, 50 μM and 500 μM aldicarb exposed L4+1 synchronized nematodes after 24 hours of intoxication. Data are shown as mean ± SEM of seven worms in four independent experiments.

### 3.3. Pharyngeal microcircuits are more sensitive to irreversible acetylcholinesterase inhibitors than to the carbamate aldicarb

In order to test the effect of irreversible organophosphate anti-cholinesterase inhibitors in the pharynx of *C. elegans*, concentration-time dependent curves were generated for pumping rate in the presence of paraoxon-ethyl, paraoxon-methyl and diisopropylfluorophosphate (DFP) (Fig. 5). Paraoxon-ethyl and -methyl were used as representative of organophosphate pesticides. In mammals, both compounds exhibit similar acetylcholinesterase inhibition rate constants (Worek, Thiermann et al. 2004). In contrast, the spontaneous hydrolysis and aging constants for them are distinct, indicating paraoxon-methyl is more likely to age the enzyme after inducing inhibition (Worek, Thiermann et al. 2004). DFP was used as representative of organophosphate nerve agents. DFP exhibits a higher inhibition and lower aging constant of mammal acetylcholinesterase compared to the paraoxon derivatives. Unlike these compounds, DFP does not show spontaneous hydrolysis, facilitating its use as a nerve agent (Worek, Thiermann et al. 2004).

**Figure 5.**
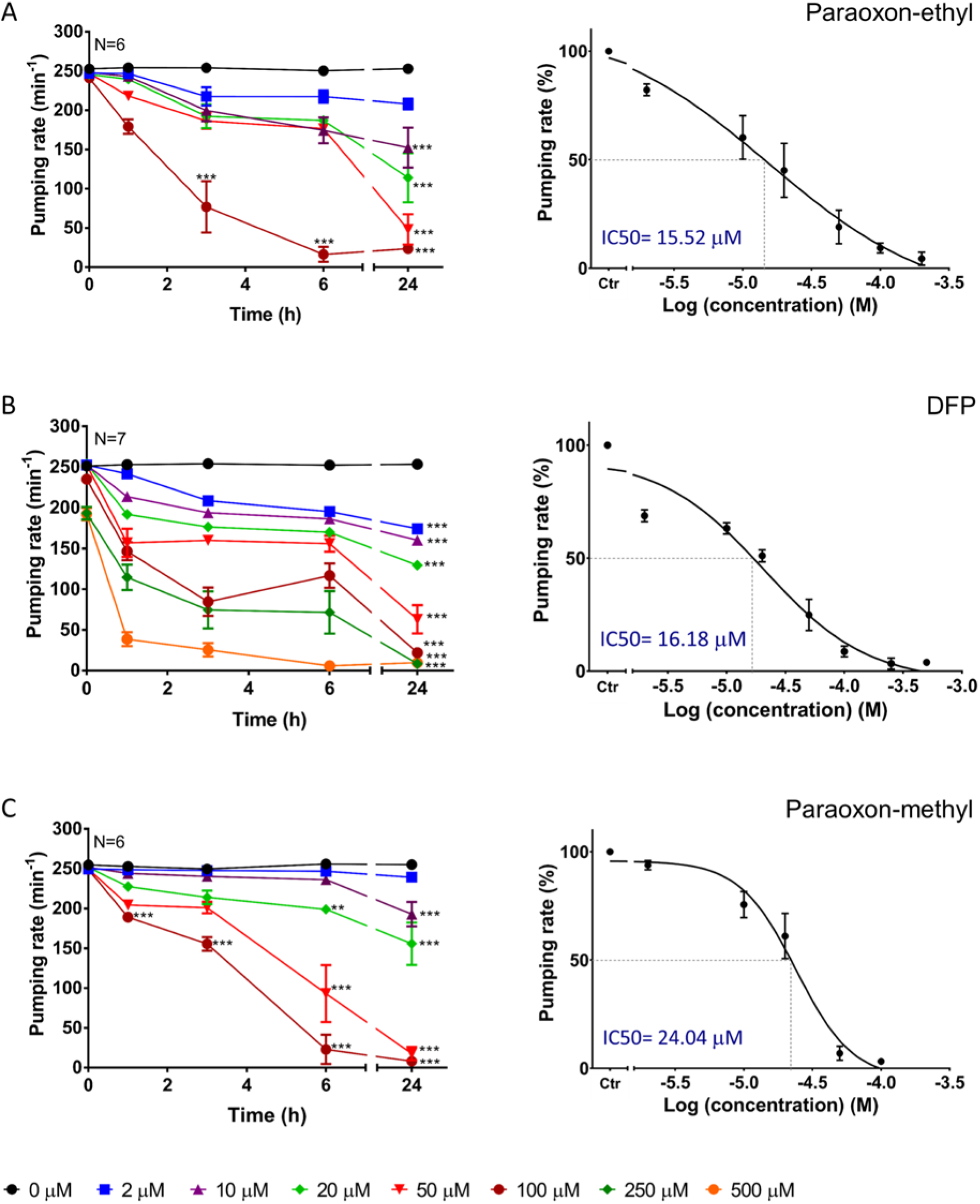
Pharyngeal pumping of *C. elegans* exposed to organophosphates. Pharyngeal pumping rate was quantified at indicated times with a range of concentrations of paraoxon-ethyl (A), DFP (B) and paraoxon-methyl (C). The IC50 values were calculated from the pump rate recorded at 24 hours of exposure to drug relative to untreated vehicle control. Data are shown as mean ± SEM of 6/7 worms in four independent experiments. *p<0.05; **p<0.01; ***p<0.001 by two-way ANOVA test.

The potency of all the organophosphates to inhibit pharyngeal pumping (Fig. 5) was much greater than observed with the carbamate (Fig. 2 and 3). The potency of inhibition of pharyngeal pumping was similar for each organophosphate. The estimated IC50 values for paraoxon-ethyl, DFP and paraoxon-methyl were 15.52 μM, 16.18 μM and 24.04 μM, respectively (Fig. 5A, 5B and 5C).

### 3.4. *C. elegans* acetylcholinesterase activity is reduced by the presence of organophosphate compounds

To characterise the inhibition of *C. elegans* acetylcholinesterase by the paraoxon derivatives and DFP, organophosphates were added to untreated worm protein. To benchmark this approach, parallel experiments using mouse brain homogenates were run. Acetylcholinesterase activity from the protein homogenates was quantified after the exposure of a single concentration of the organophosphate at increasing times of incubation (Fig. 6).

**Figure 6.**
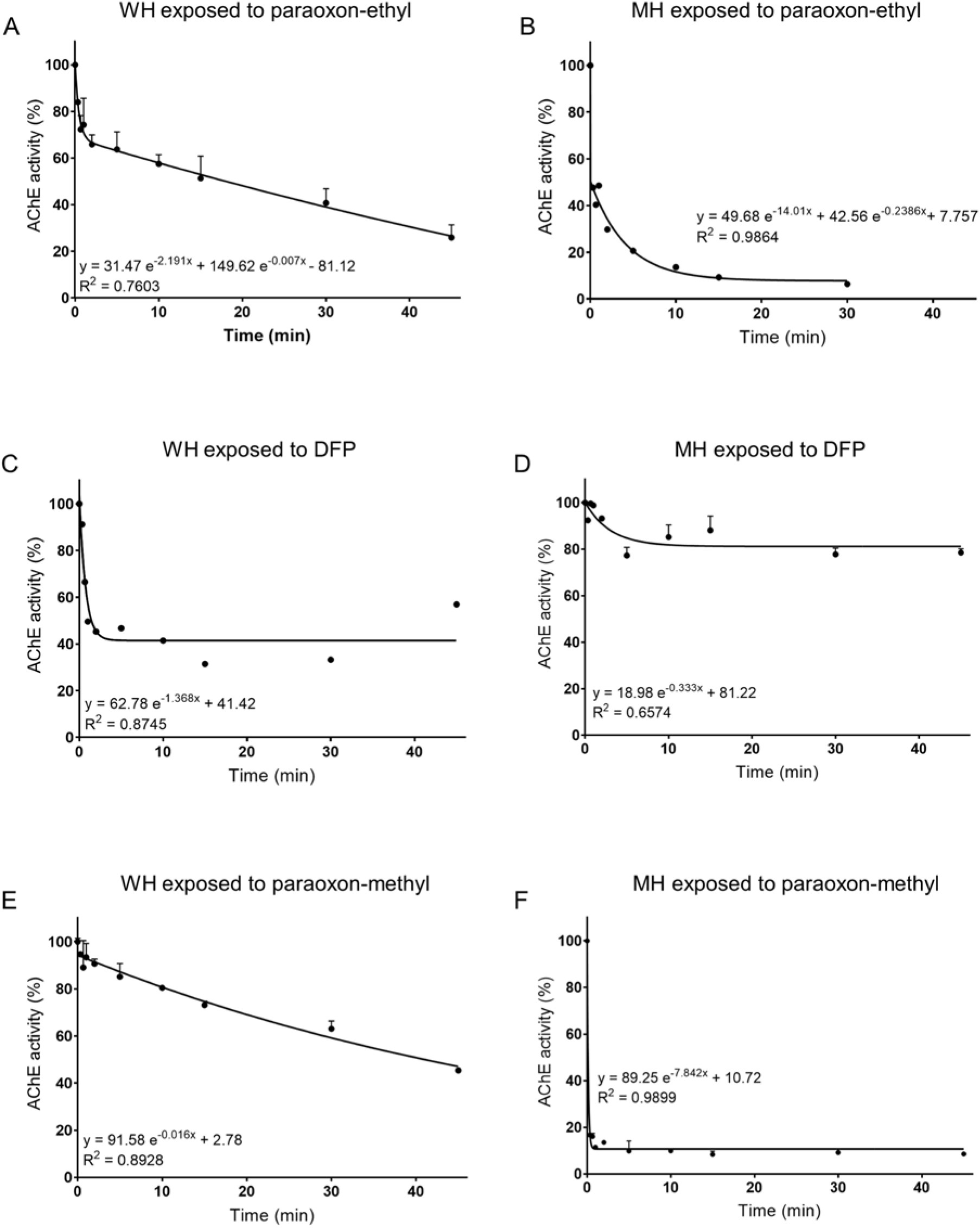
Paraoxon-ethyl, DFP or paraoxon-methyl show a time dependent inhibition of the acetylcholinesterase activity associated with *C. elegans* and mouse brain homogenates. Worm (WH) and mouse brain (MH) homogenates were incubated in addition of 1 μM of paraoxon-ethyl, DFP or paraoxon-methyl to allow timed incubation of the enzyme inactivation before synchronized measurement of homogenate associated acetylcholinesterase activity. Acetylcholinesterase activity was expressed as percentage of the unexposed homogenate activity. Two-phase exponential decay curve was ascribed as the best fit for the inhibition of nematode (A) and mouse (B) acetylcholinesterase activity at different end-point times of incubation with paraoxon-ethyl. Single exponential decay curve was fitted to the inhibition of worm (C) and mouse (D) acetylcholinesterase with DFP. One-phase inhibition of worm (E) and mouse (F) acetylcholinesterase inhibition with paraoxon-methyl at different end-point times of incubation with the organophosphate.

Paraoxon-ethyl inhibition over time showed two different phases in both the *C. elegans* and mouse homogenate (Fig. 6A and 6B). The inhibition rate was greater at the first times of intoxication than in later exposure times.

Upon incubation with DFP, both the nematode and mouse acetylcholinesterase activity was diminished within the first 2 minutes of exposure, after which it levelled off to a steady state inhibition (Fig. 6C and 6D). However, the steady state inhibition of worm acetylcholinesterase reached after 2 minutes of incubation with DFP was about 60% of the total activity while the inhibition of mouse acetylcholinesterase was only 20%. It might indicate that *C. elegans* acetylcholinesterase is more susceptible for DFP inhibition than the mouse enzyme in the conditions assayed.

Paraoxon-methyl inhibition over the time in *C. elegans* and mouse homogenate fitted a single decay curve. Nevertheless, the inhibition of worm acetylcholinesterase is progressive over the time while the reduction of the mouse acetylcholinesterase activity is more rapid, reaching steady state within 1 min of incubation (Fig. 6E and 6F).

Overall, the reduction of the pharyngeal pumping rate in the presence of the anti-cholinesterases was associated with a time and concentration dependent inhibition of the nematode acetylcholinesterase. Based on the degree of organophosphate-induced inhibition of behaviour and homogenate associated acetylcholinesterase activity, the results suggest that DFP is more efficient than paraoxon-ethyl, which is more efficient than paraoxon-methyl, even if the IC50 values and homogenate inhibition reached similar values at the longest incubation time (Fig. 5 and 6).

### 3.5. Recovery of pharyngeal function from organophosphates intoxication

To investigate if pharyngeal pumping recovered from anti-cholinesterase intoxication, nematodes were incubated with inhibitors for 24 hours. The intoxicated nematodes were then transferred onto control plates, or ones treated with obidoxime or pralidoxime. These oximes, which did not affect pumping themselves, were investigated to see if their known ability to facilitate recovery is manifest in *C. elegans* (Fig. 7).

**Figure 7.**
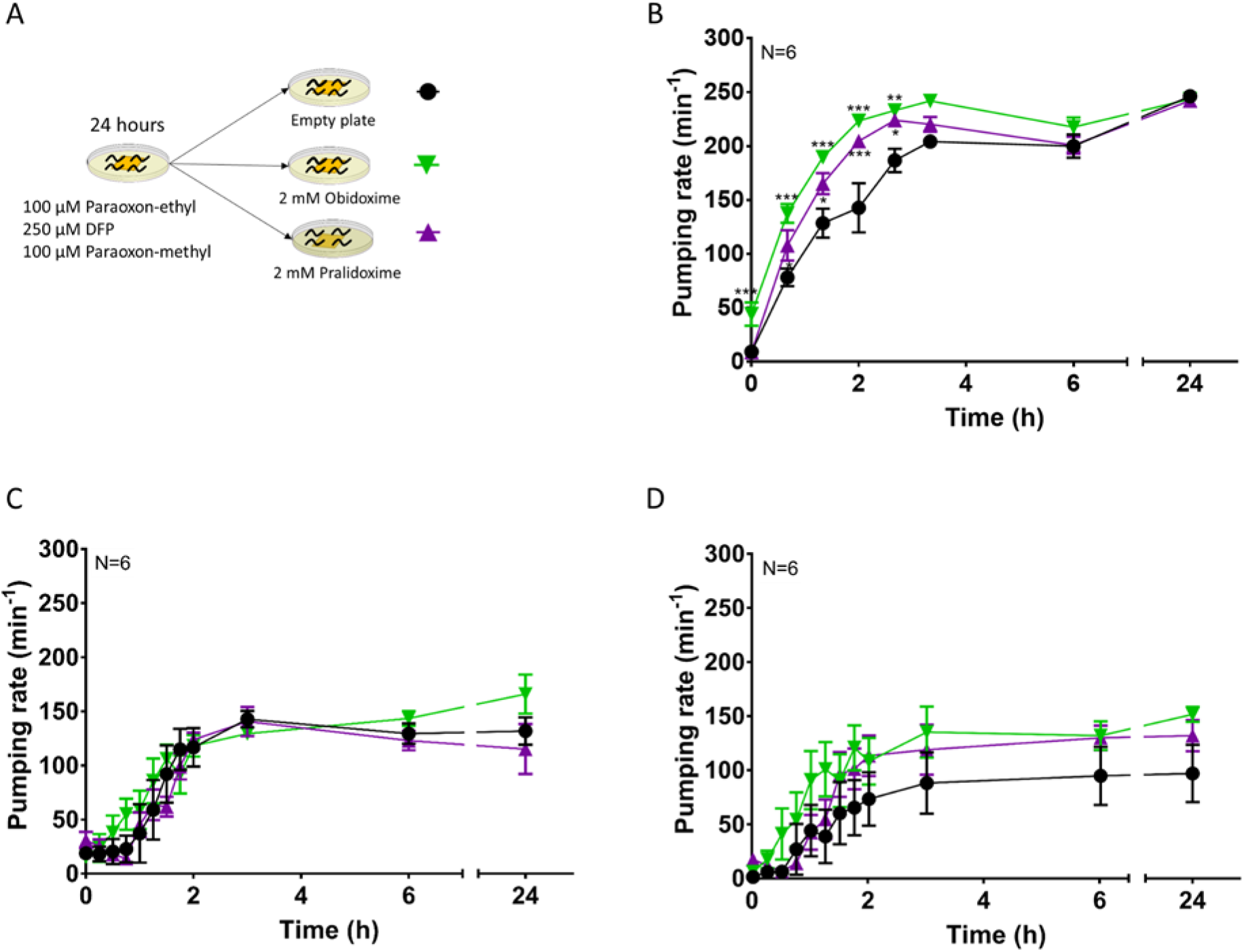
Spontaneous and oxime induced recovery of pharyngeal pumping inhibition from organophosphates intoxication. A) Experimental design of pharyngeal function recovery after 100 μM paraoxon-ethyl, 250 μM DFP or 100 μM paraoxon-methyl intoxication. Synchronized L4+1 nematodes were incubated on drug-containing plates for 24 hours. After transfer to control, obidoxime or pralidoxime containing plates pumping was scored at indicated times. B) Nematodes intoxicated on paraoxon-ethyl plates exhibited a fast and complete recovery of pumping enhanced by oxime treatment. C) Nematodes intoxicated on DFP plates for 24 hours displayed slow, incomplete and oxime-insensitive recovery. D) Recovery from paraoxon-methyl was incomplete and oxime-independent. Data are shown as mean ± SEM of six worms in six independent experiments. *p<0.05; **p<0.01; ***p<0.001 by two-way ANOVA test.

Nematodes intoxicated with 100 μM paraoxon-ethyl exhibited a fast recovery of the pharyngeal function, which was complete at 4 hours after being transferred onto empty plates (Fig. 7B). Recovery was accelerated when nematodes were removed from organophosphate and placed onto either the obidoxime or the pralidoxime plates. Obidoxime was the more effective of the two compounds tested in rescuing the pharyngeal activity, with a half-time of recovery of 1 h compared to 1.33 h on pralidoxime or 2 h on control plates (Fig. 7B). In contrast, nematodes incubated on either 100 μM paraoxon-methyl or 250 μM DFP plates for 24 hours did not show complete recovery when transferred to drug free plates (Fig. 7C and 7D). Moreover, the exposure of intoxicated worms to either of the oximes tested did not improve the rescue of the pharyngeal function.

Overall, the complete, fast and oxime-sensitive recovery of the pharyngeal function after paraoxon-ethyl exposure indicates that paraoxon-ethyl is not able to age the worm acetylcholinesterase and the presence of oximes facilitates the reactivation of the inhibited enzyme. In contrast, the slow, incomplete and oxime-insensitive recovery of the pumping rate after either paraoxon-methyl or DFP intoxication suggests these compounds irreversibly inhibit *C. elegans* acetylcholinesterase.

### 3.6. Nematode acetylcholinesterase recovery after organophosphate inhibition

To test if the organophosphates that inhibit pharyngeal pumping irreversibly modify the worm acetylcholinesterase, we investigated the recovery of the organophosphate-inhibited homogenates with and without the post intoxication addition of obidoxime.

Both worm and mouse acetylcholinesterase inhibited by paraoxon-ethyl had a significant but incomplete recovery by obidoxime (Fig. 8A and 8B).

**Figure 8.**
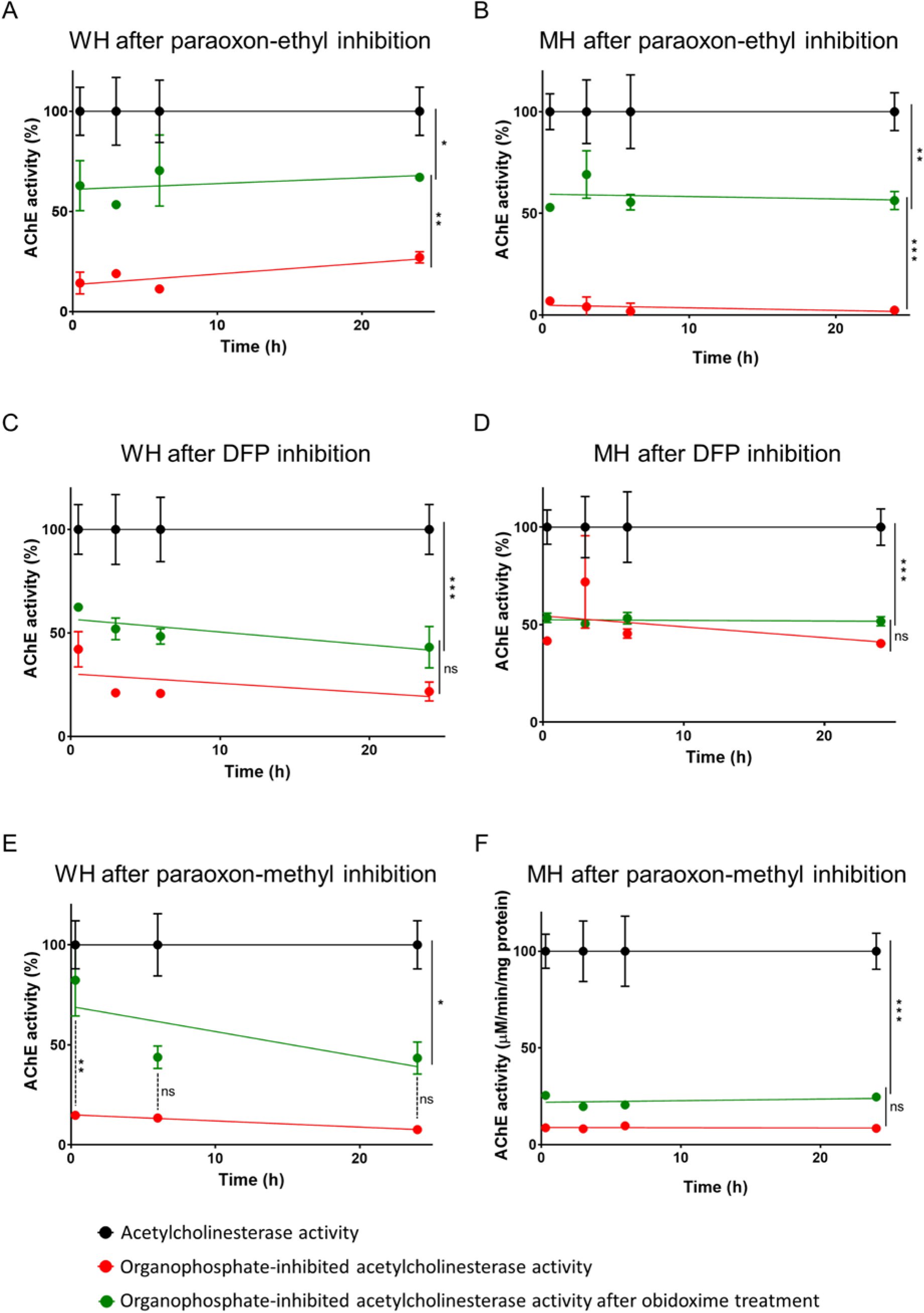
*C. elegans* and mouse acetylcholinesterase is aged after the inhibition with either DFP or paraoxon-methyl. OP-inhibited worm (WH) or mouse (MH) acetylcholinesterase activity was quantified in the presence or absence of a single concentration of obidoxime. Untreated homogenates were used as controls. Acetylcholinesterase activities were represented as the percentage of activity referred to the untreated control. A) Worm homogenate inhibited by paraoxon-ethyl exhibited an 80% reduction of the acetylcholinesterase activity that was partially recovered by the obidoxime treatment. B) Mouse homogenate acetylcholinesterase inhibited by paraoxon-ethyl displayed a partial recovery of its activity after obidoxime treatment. C) The obidoxime treatment did not significantly improved the acetylcholinesterase activity of the worm acetylcholinesterase inhibited by DFP. D) Mouse acetylcholinesterase activity was reduced after DFP treatment in 50%. Nevertheless, there is not recovery of the enzyme activity after the obidoxime treatment. E) Worm homogenate exposed to paraoxon-methyl displayed a reduction of the acetylcholinesterase activity that can be recovered by the obidoxime treatment after 30 min of incubation. However, there is not recovery in the consequent end-point times tested. F) The inhibition of mouse acetylcholinesterase activity by paraoxon-methyl was not recovered by the obidoxime treatment. *p<0.05; **p<0.01; ***p<0.001 by two-way ANOVA test.

As already observed (Fig. 6C and 6D), the inhibition of worm acetylcholinesterase by DFP was greater than the inhibition of mouse acetylcholinesterase (Fig. 8C and 8D). After the incubation step with obidoxime, there was no significant recovery of the acetylcholinesterase activity supporting enzyme inhibition by this organophosphate had progressed through an irreversible reaction.

Finally, worm/mouse acetylcholinesterase exposed to paraoxon-methyl exhibited a nearly complete reduction of the activity (Fig. 8E and 8F). The presence of obidoxime did not improve the acetylcholinesterase activity of the mouse paraoxon-methyl enzyme indicating a rapid irreversible inhibition (Fig. 8F). Nonetheless, the effect of obidoxime in the worm paraoxon-methyl inhibited acetylcholinesterase indicated a time-dependent process. At 30 min after removing the excess of organophosphate, the presence of obidoxime recovered the total acetylcholinesterase activity, indicating a reversible reaction of inhibition at this time point. At subsequent times beyond 30 min, there was no improvement in the acetylcholinesterase activity of paraoxon-methyl inhibited enzyme by the incubation with obidoxime (Fig. 8E). This is consistent with a progressed inhibition through a classic aging reaction (Sun, Chang et al. 1979) or alternatively an inability of oxime to execute a nucleophilic attack at the organophosphate bound to the serine at the active site (Worek, Thiermann et al. 2004, Worek, Thiermann et al. 2016).

## 4. Discussion

### 4.1. Pharyngeal pumping rate as mechanism for evaluating the effect of anti-cholinesterase intoxication

In the present study, we have used whole organism intoxication of *C. elegans* to investigate carbamate and organophosphate poisoning of cholinesterase. The study verifies previous results that worm behaviours are dependent on cholinergic transmission and therefore suitable to investigate anti-cholinesterase intoxication (Cole, Anderson et al. 2004, Boyd, McBride et al. 2007, Rajini, Melstrom et al. 2008, McVey, Mink et al. 2012, Leelaja and Rajini 2013). Most of the previous studies were focused on the direct effect of anti-cholinesterases on *C. elegans* movement, either in liquid or solid culture, while the pharyngeal effect has been indirectly scored by the reduction of food availability in liquid culture (Boyd, McBride et al. 2007, Rajini, Melstrom et al. 2008, Boyd, Smith et al. 2010). However, the recovery of those behaviours after acetylcholinesterase inhibitors exposure has never been probed. This is an important area for investigating mitigation approaches and cross-referencing to the similarity of core mode of action in the model organisms and humans.

The pharyngeal pump depends on acetylcholine excitation of the pharyngeal muscles to drive the contraction and relaxation cycle that allows the food intake (Avery 1993, McKay, Raizen et al. 2004, Boyd, McBride et al. 2007). This readily scored behaviour on plates offers a distinct route to test acetylcholinesterase intoxication and recovery. Comparing the sensitivity of the pump rate to intoxication relative to the shrinkage of the worm or the binary scoring of paralysis suggests this assay may be more sensitive and better suited to discern the incremental concentration-dependent and recovery effects. The assay, which is conducted on the worms on food, has a good dynamic range. Pumping is elevated from about 40 to 250 pumps per minute when worms enter the food and the concentration-dependent inhibition of this activity by the anti-cholinesterase appears to operate across this dynamic range. Indeed, when judged as an observer based bio-assay, it is more sensitive than locomotion, previously described as a phenotype for assessing organophosphate intoxication (Cole, Anderson et al. 2004, Melstrom and Williams 2007, Leelaja and Rajini 2013). The pharyngeal neuromuscular innervation of *C. elegans* consists of a subset of cholinergic and glutamatergic neurons that synapse onto the radial muscles of the pharynx (Albertson and Thomson 1976, Trojanowski, Raizen et al. 2016). The release of acetylcholine mainly by the MC and M4 motor neurons results in the contraction of the muscles causing the opening of the lumen and therefore the entering of bacteria (Albertson and Thomson 1976, Trojanowski, Raizen et al. 2016). In the presence of anti-cholinergic compounds, the pharyngeal muscles remain hypercontracted and the lumen continuously open (Fig. 2B) causing the paralysis of the pharyngeal movement (Fig. 2A). We demonstrated that the reduction of the pumping rate was better correlated with the inhibition of the acetylcholinesterase activity by aldicarb compared to body wall neuromuscular junction phenotypes (Fig. 4). This fact might be due to a differential sensitivity of the pharyngeal circuits to intoxication with acetylcholinesterase inhibitors compared to body wall circuits. The ingestion of the anti-cholinesterase compounds with the bacteria while feeding might be a faster access pathway for the inhibitors than throughout the cuticle. However, the pharyngeal movement quantification is also a better discriminatory assay, ranging the impact of intoxication from 0 to 250 pumps/min while there is not such an incremental effect in the paralysis of the locomotion.

The intoxication of the pharyngeal muscles by organophosphates caused a reduction of the pumping rate, which gave the rank order of potency of toxicity: paraoxon-ethyl > DFP > paraoxon-methyl with slight differences of the IC50 values between them (Fig. 5). Similar to mammalian investigations, the acute toxicity of organophosphates was associated with the block of the acetylcholinesterase activity by the inhibitors (Fig. 6). In fact, the biochemical reduction of acetylcholinesterase activity during exposure indicates a similar ranking of toxicity as the one measured with pharyngeal pumping (Fig. 5, 6A, 6C, 6E). It might indicate that organophosphates can easily access the worm acetylcholinesterases when the nematodes are on inhibitor-containing plates. They block enzyme activity, causing the hypercontraction of the pharyngeal muscles and therefore the paralysis of the feeding. The route of the drug access into the worm is still unknown; it might be through the cuticle but also by ingestion when they feed.

The action of organophosphates in the mouse homogenate indicates that acetylcholinesterase is more susceptible to the inhibition by either paraoxon-ethyl or -methyl than by DFP (Fig. 6B, 6D, 6F). This is consistent with acute toxicity and kinetic data previously published for murine models poisoned with organophosphates where LD50 values and inhibition constants for DFP are slightly higher than for paraoxon-ethyl or -methyl, independently of the mode of administration (Johnson and Wallace 1987, Gearhart, Jepson et al. 1990, Misik, Pavlikova et al. 2015).

The different rank of toxicity for DFP between *C. elegans* and mouse acetylcholinesterase might indicate a difference in the kinetics of inhibition by the OP between the two organisms. This variance has been previously described among the diverse organism models probed for their reactivity acetylcholinesterase inhibition and recovery (Johnson and Wallace 1987, Gearhart, Jepson et al. 1990, Worek, Thiermann et al. 2004, Worek, Aurbek et al. 2008, Coban, Carr et al. 2016). Despite the different rank of toxicity, both *C. elegans* and mouse DFP-inhibited acetylcholinesterase exhibited no recovery after obidoxime treatment (Fig. 8C and 8D), which is consistent with the absence of recovery observed in the pharyngeal pumping after DFP intoxication (Fig. 7C).

### 4.2. Pharyngeal pumping rate as a metric for evaluating spontaneous recovery and reactivation after organophosphate intoxication

The recovery of the acetylcholinesterase activity is key to treat the cholinergic syndrome. Oxime treatment in humans after OP poisoning offers an established supporting therapy (Eddleston and Chowdhury 2016). However, the recovery and the oxime efficiency is an OP-dependent process (Worek, Thiermann et al. 2004). In mammalian models, the rate of the reaction and the efficiency of possible therapies can be analysed biochemically by quantifying the acetylcholinesterase activity either in blood or brain samples of intoxicated animals (Maxwell, Brecht et al. 1987, Bajgar 1992, Misik, Pavlikova et al. 2015). We describe here a simple *in vivo* experiment in a model organism that is potentially indicative of the chemical state of the acetylcholinesterase active site after the organophosphate intoxication. The recovery of the pharyngeal function after paraoxon-ethyl exposure and the improvement by the oximes (Fig. 7B) supports an oxime-sensitive reaction between the OP and the worm acetylcholinesterase (Fig. 8A). In biochemical experiments using worm protein, we observed a reduction of acetylcholinesterase activity that can be rescued by the incubation with obidoxime (Fig. 8A). In contrast, the inefficiency of the oxime treatment after paraoxon-methyl or DFP inhibition indicates an irreversible modification of the nematode enzyme by these organophosphates (Fig. 8C and 8E). This was manifest by the failure to recover pharyngeal function when intoxicated worms are transferred onto either empty or oxime-containing plates (Fig. 7C and 7D).

The biochemical study of spontaneous recovery and oxime-sensitive reaction in mouse acetylcholinesterase is consistent with previously published data (Tripathi and Dewey 1989, Kardos and Sultatos 2000). Non-aged acetylcholinesterase after the exposure of paraoxon-ethyl was able to recover partially the activity in the presence of obidoxime (Fig. 8B) while the aged acetylcholinesterase that predominates after either paraoxon-methyl or DFP inhibition could not be rescued by the oxime treatment (Fig. 8D and 8F) (Tripathi and Dewey 1989, Kardos and Sultatos 2000).

To conclude, the analysis of the nematode pharyngeal function after OP intoxication might be indicative of the acetylcholinesterase state after the enzyme inhibition, spontaneous and obidoxime-induced reactivation.

## 5. Conclusion

In previous studies, *C. elegans* body wall phenotypes have been used to understand acetylcholinesterase inhibition by organophosphate exposure and, in some of them; it was correlated with the quantification of acetylcholinesterase activity in the worm (Melstrom and Williams 2007, Rajini, Melstrom et al. 2008, Leelaja and Rajini 2013). Here, we demonstrated in the present study that the pharyngeal function represents a more precise phenotype to understand acetylcholinesterase inhibition by OP drugs. Interestingly, the rescue of the phenotype was also correlated with the rate of the acetylcholinesterase reaction upon OP inhibition as well as the efficiency of the reactivators. It makes the pharynx of *C. elegans* an attractive tool for discovering new drugs able to reactivate the inhibited acetylcholinesterase. Furthermore, clear benchmarking of this class of neurotoxicological agents in a tractable bio-assay in *C. elegans* means that genetic manipulation of these effects can be probed. This provides a new approach to investigate mitigation of such neurotoxicity that may translate to human poisoning.

## Acknowledgements

We thank Aleksandra Pitera and Dr. Katrin Deinhardt for providing mouse brain homogenate. Additionally, *C. elegans* strains were provided by the CGC, which is funded by NIH Office of Research Infrastructure Programs (P40 OD010440).

## Funding

This work was equally funded by the University of Southampton (United Kingdom) and the Defence Science and Technology Laboratory, Porton Down, Wiltshire (United Kingdom).

## Author contributions

**Patricia G. Izquierdo**: Conceptualization, Data curation, Formal analysis, Investigation, Methodology, Validation, Visualization, Roles/Writing - original draft. **Vincent O’Connor**: Conceptualization, Funding acquisition, Methodology, Supervision, Writing - review & editing. **Christopher Green**: Conceptualization, Funding acquisition, Methodology, Supervision, Writing - review & editing. **Lindy Holden-Dye**: Conceptualization, Funding acquisition, Methodology, Supervision, Writing - review & editing. **John Tattersall**: Conceptualization, Funding acquisition, Methodology, Supervision, Writing - review & editing.

---

**Supplementary figure 1.**
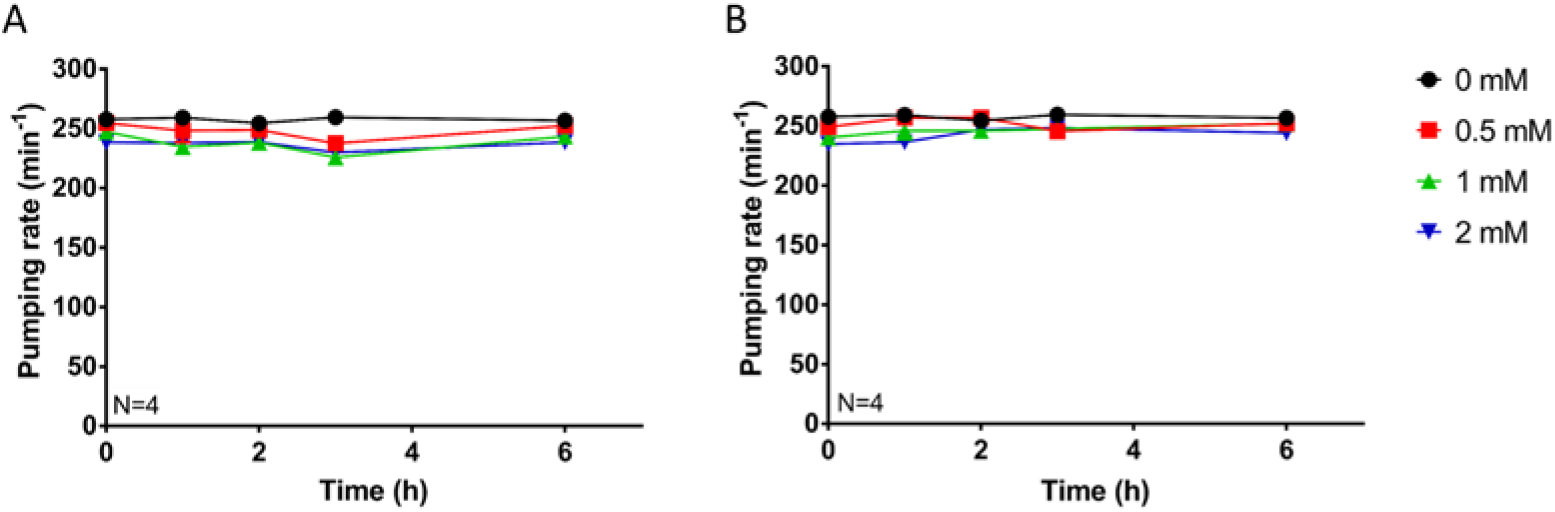
Pharyngeal pumping rate phenotype of *C. elegans* wild type adults exposed to oximes plates. Pharyngeal pumping rate per minute was quantified at different end-point times for synchronized L4+1 nematodes exposed to increasing concentration of plates containing obidoxime (A) or pralidoxime (B). Neither obidoxime nor pralidoxime had any effect in the pharyngeal function of *C. elegans*. Data are shown as mean ± SEM of four worms in four independent experiments.

## Notes

### Competing Interest Statement

The authors have declared no competing interest.

